# A robot model of compass cue calibration in the insect brain

**DOI:** 10.64898/2026.06.25.734539

**Authors:** Robert Mitchell, Marie Dacke, Barbara Webb

## Abstract

Dung beetles can use a variety of orientation cues to maintain a consistent bearing during ball-rolling. Where several cues are available, they appear to learn the spatial relationship between them, providing redundancy if some cues are removed. Mounting evidence indicates that such a learning process is implemented in the insect head direction circuit; specifically, in the plastic substrate between sensory input neurons and compass neurons in the central complex. This plasticity appears to be driven by rotational movements, providing a clear link with observed beetle ‘dance’ behaviour. Here, we extend our functional model of this circuit and use it on a robot platform, to test it in the same behavioural assay as was used for the beetles. The robot was able to replicate the beetle’s ability to substitute a directional wind cue for a point source light cue in guiding straight-line movement. However, it also revealed significant biasing coupled to dance direction. This biasing appears to be caused by inherent conflict between recurrent and instantaneous inputs to the compass circuit. We predict that the real insect should experience similar issues unless it has evolved a neural mechanism to compensate.

## 1 Introduction

Insects (along with many other animals [7]) are known to integrate multiple cues to guide their navigation behaviours. In order to combine different sensory modalities, the information must be associated to the same frame of reference to produce coherent rather than conflicting estimates. This also ensures that animals can generate consistent behaviour when using any cue in isolation and is particularly useful in cases where one cue is intermittently available or perhaps subject to change (either with respect to other cues or the underlying property the animal is trying to estimate).

For example, bushcrickets can learn the relationship between an intermittent auditory cue (the calling song of a potential mate) and a visual scene [48], allowing them to continue moving in the correct direction during gaps in the song. Bogong moths appear to learn the relationship between starlight patterns and magnetic information [14], allowing them to navigate consistently, despite the apparent change in star patterns over the duration of their migration. Similarly, ball-rolling dung beetles are able to learn arbitrary cue arrangements, even where the arrangement (e.g. the relationship of sky polarisation direction to sun location) should never occur in their natural surroundings [20].

In insects, compass cue calibration has been proposed to occur in the central complex (CX; a set of midline neuropils conserved across all insects studied to date [30]), specifically in the plastic interface between sensory input (ER) neurons and compass (EPG) neurons [38, 21, 51, 35]. For each modality, all input neurons connect to all compass neurons and these connections can be remapped so as to bring different cue inputs into the same frame of reference (figure 1b). Despite the acceptance of this conceptual model [21, 51], it is difficult to find behavioural examples which can be concretely coupled to this mechanism.

**Figure 1.**
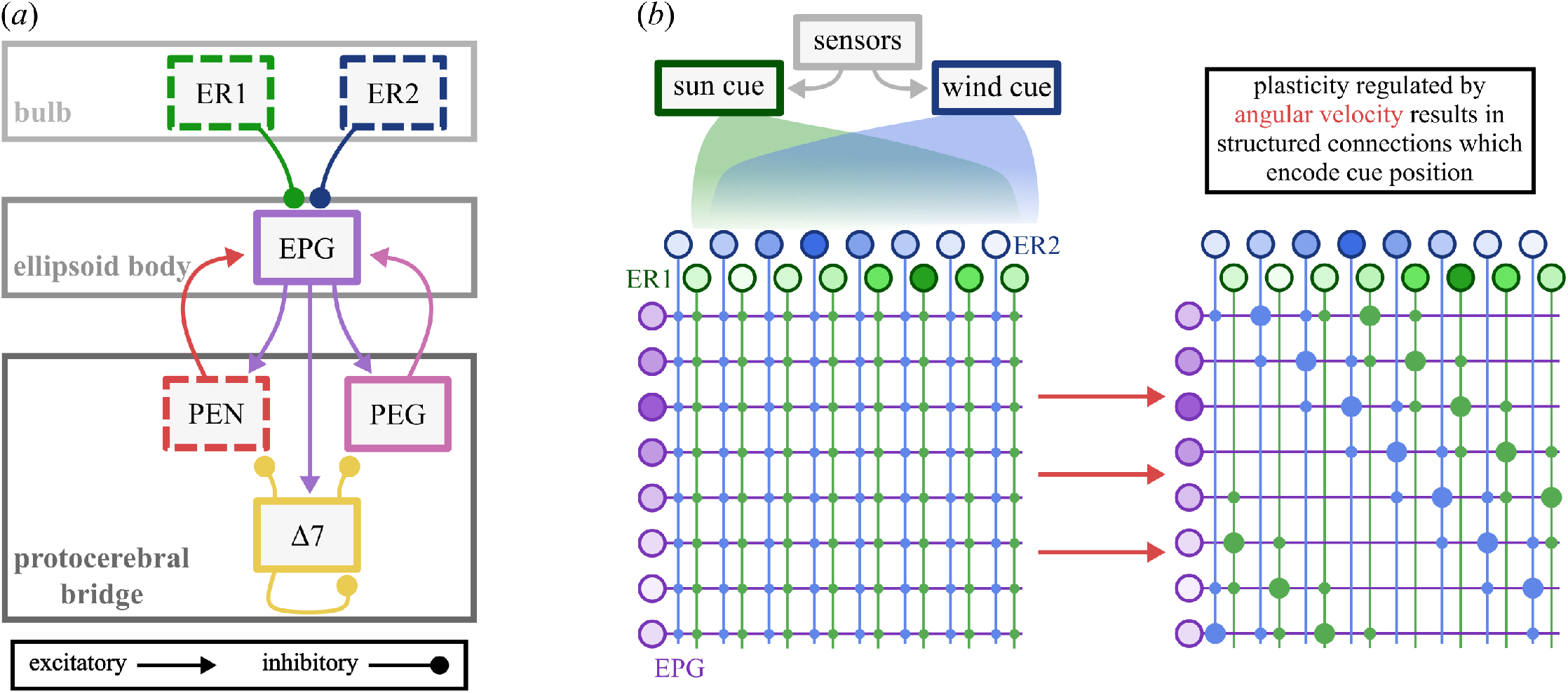
Model overview. (*a*) The neural model from Mitchell et al. [35] is made up of six neural populations. Dashed borders indicate model inputs. **ER** neurons act as cue inputs (one population for each cue). **EPG**s are the “compass” neurons which encode the agent’s orientation as a sinusoidal activity bump. **PEN**s act as angular velocity inputs to the compass. **PEG**s provide recurrent circuitry which helps to maintain network activity in the absence of external input. Δ7s provide lateral inhibition which helps to stabilise the compass bump. (*b*) The cue calibration mechanism with which this paper is concerned. All ER neurons connect to all EPGs via plastic connections. As the agent rotates, these connections are remapped in an anti-Hebbian fashion which (in simulation) results in structured connections which encode the spatial relationship of the different cues. This calibration allows an agent to orientate consistently regardless of which cue it is using [21, 38].

One clear behavioural example can be found in ball-rolling dung beetles performing straight-line orientation [11]. After arriving at a dung source, the beetles will cut off a piece of dung, form it into a ball, perform a brief orientation dance, and then roll it away while maintaining a straight course.

The beetles learn the spatial relationship of the available orientation cues [10, 20], and the moment this learning occurs appears to be the orientation dance [20]. This is compelling as the dance is a rotational behaviour [1], which aligns with the proposed neural substrate (where learning is coupled to rotation). Moreover, the orientation cues available to the beetles (sun [5], moon [9], polarised light [16], wide-field intensity [16], spectral information [17, 52], and wind [10]) all appear to have a pathway to the compass via ER neurons (visual [42, 18], polarisation [29, 18], and wind [47, 38]).

In summary, the beetles’ orientation cues have pathways into the proposed CX substrate and the learning is behaviourally linked to rotation, as is true within the CX substrate [22]. Dung beetles therefore provide a behavioural example which can be firmly linked to this neural mechanism.

We have developed a computational model of the multimodal insect compass, and shown that it can produce cue calibration behaviour in simulated dung beetles [35]. Here, we extend this model by updating the learning mechanism to operate flexibly in proportion to angular velocity [22], and to include noisy neurons. We also add a bio-inspired steering circuit, based on the work of Mussells Pires et al. [36] and Mitchell and Webb [34]. The updated neural model is deployed on a custom robot platform to allow us to test the model as we would an insect, using real sensory cues in an experimental paradigm identical to that used for dung beetles [10]. Our robot experiment confirms that the updated model does work as we expect, but exposes several problems with our high-level understanding of this circuit which have not been raised by previous works.

## 2 Methods

### 2.1 Neural model overview

Our neural model was originally designed to investigate cue integration in the insect compass (of which cue calibration was one component). Details of the model can be found in Mitchell et al. [35] (or in our electronic supplementary information), but a brief overview is given here and in figure 1. The core of the neural circuit is a ring attractor formed by the EPG (“compass”) neurons in the ellipsoid body, which are coupled to each other by excitatory (PEG) and inhibitory (Δ7) feedback to maintain a single activity bump. Offset feedback connections (PEN) modulated by self-motion can rotate the bump around the ring. The EPG neurons also get input from external sensory cues via ER neurons (note that we use the updated nomenclature of ‘ER’ neurons for ring neurons, the same neurons are termed ‘R’ neurons in [35]). The connections from ER to EPG neurons are inhibitory and plastic; an anti-Hebbian learning rule weakens the connection between simultaneously active ER and EPG neurons, reducing the inhibition and thus calibrating the external cue’s input to the circuit with the current compass bump.

### 2.2 Model updates

#### 2.2.1 Modifications to the compass circuit

##### Flexible learning rate

In the original model, learning between ER and EPG neurons was either disabled or enabled at a set rate during specific learning events [35]. Recent work has shown that plasticity between ER and EPG neurons is regulated flexibly by angular velocity [22]. Thus, the learning rate *η* is now set to be proportional to angular velocity:

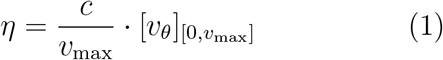

where *c* is a tuning constant, *v*_max_ is the maximum angular velocity which can be perceived by the network via PEN inputs (see [35]), and *v*_*θ*_ is the instantaneous absolute angular velocity. [*x*]_[*y,z*]_ = max(*y*, min(*x, z*)) is a clipping operation. The learning rule itself is identical to [35], only the rate parameter is changed:

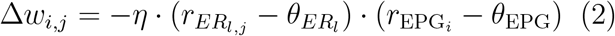

where Δ*w*_*i,j*_ is the change in synaptic strength from ER_*l,j*_ onto EPG_*i*_, the *r* terms are spiking rates for the respective neurons and the *θ* terms are activity thresholds. Learning is continuously enabled.

##### Flexible ER neuron regulation

In [35], ER neuron input to EPGs was reduced (but not eliminated) during learning events. As learning rate is now proportional to angular velocity, so too is this reduction:

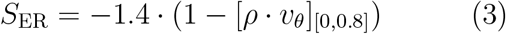

where *v*_*θ*_ is the absolute instantaneous angular velocity and *ρ* is a tuning constant. ER neurons are inhibitory and, in the model, *™*1.4 is their basic scalar weight onto EPGs; |*S*_ER_| reduces as angular velocity increases.

This modulates the relative strength of compass inputs during rotation. As angular velocity increases, ER input to EPGs is down-regulated and relative influence from PENs and PEGs rises. When angular velocity decreases, ER neurons will have more effect. This essentially means that proprioceptive inputs drive the compass during moments of high angular velocity with allothetic inputs anchoring the compass during moments of low angular velocity.

##### Reweighting of internal feedback

Early robot testing showed that the original model was brittle to externally imposed movement of a short duration, such as might occur naturally if, for example, a beetle was displaced by a wind gust (or an experimenter). Moving the robot by hand could cause neural activity to die off completely, meaning no directional bump was maintained. This had not been noticed in the simulation environment in which the model was developed, as the simulated displacements were instantaneous. The network has therefore been re-tuned to improve robustness. Practically this amounts to a small up-weighting of the protocerebral bridge inputs to the EPGs (the PENs and PEGs), compared to the original implementation [35]. Increasing the influence of PEGs corresponds to an up-regulation of the internal feedback that maintains the bump.

##### Neural noise

Neurons were noiseless in the original model because one of our primary aims was to investigate whether the circuitry could implement a vector calculation for cue integration under ideal conditions. For the current work, we added a small amount of Gaussian noise to the neuron activation function:

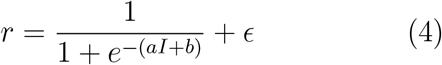

where *I* is the input, *a* and *b* give the slope and bias of the activation function, and *ϵ ∼ N* (0, 0.001).

This aids biological plausibility (as neurons are not perfect) but also has a practical function in preventing neuron activity from degenerating into a state where all activities are equal (from which it can be impossible to recover). As discussed next, it also turns out to solve the problem of network initialisation.

##### Spontaneous network initialisation

In [35], the network was rotated on the spot with allothetic and idiothetic inputs which would cause bumps of activity to form in each layer and stabilise. In this work, the addition of activation noise and the slight retuning of EPG inputs seems to cause activity bumps to appear and stabilise spontaneously, without any need for rotation. Noise causes small differences in the activities of neighbouring neurons, which get reinforced by the recurrent circuitry, eventually leading to a stable sinusoidal activity pattern.

This improves the biological plausibility of the model as the old version required either structured ER→ EPG connections to generate activity bumps, or activity bumps to generate structured connections. This chicken-and-egg problem seems to be solved by the circuit architecture reinforcing small activity imbalances generated by noisy neurons. This also explains how the network could recover in the case that an activity bump is completely lost (something the previous version could not do). Finally, it may help account for the observation (in flies) that there is a random anatomical offset of the bump, relative to external cues, between animals or on different behavioural bouts, as the initial spontaneous location of the bump will determine how it is mapped to external cues.

#### 2.2.2 Steering circuit

To control the robot’s behaviour, the compass circuit has to produce steering signals as output. We implemented a biologically inspired steering circuit that captures the steering principle described by Mussells Pires et al. [36] for goal-directed navigation in the fruit fly brain. Further discussion of this principle and the possible ways it could be instantiated in neural circuits is provided in [34].

##### FC2 (goal direction) neurons

Each FC2 neuron has a specific preferred angle, generating a sinusoidal signal which acts as a population vector encoding for a goal direction.

The input for the *i*th FC2 neuron is given by:

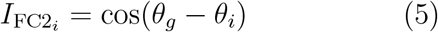

where *θ*_*g*_ is the goal direction and *θ*_*i*_ is the preferred angle of the *i*th FC2 neuron. This formulation is essentially identical to that given by [36] (they include a non-linear transformation which is fitted to their behavioural data).

##### PFL3 (steering) neurons

PFL3 neurons split into two populations which steer the agent either to the left or the right. Steering neurons form pairs (one from each population) which drive the agent along a particular bearing with respect to the compass (figure 2).

**Figure 2.**
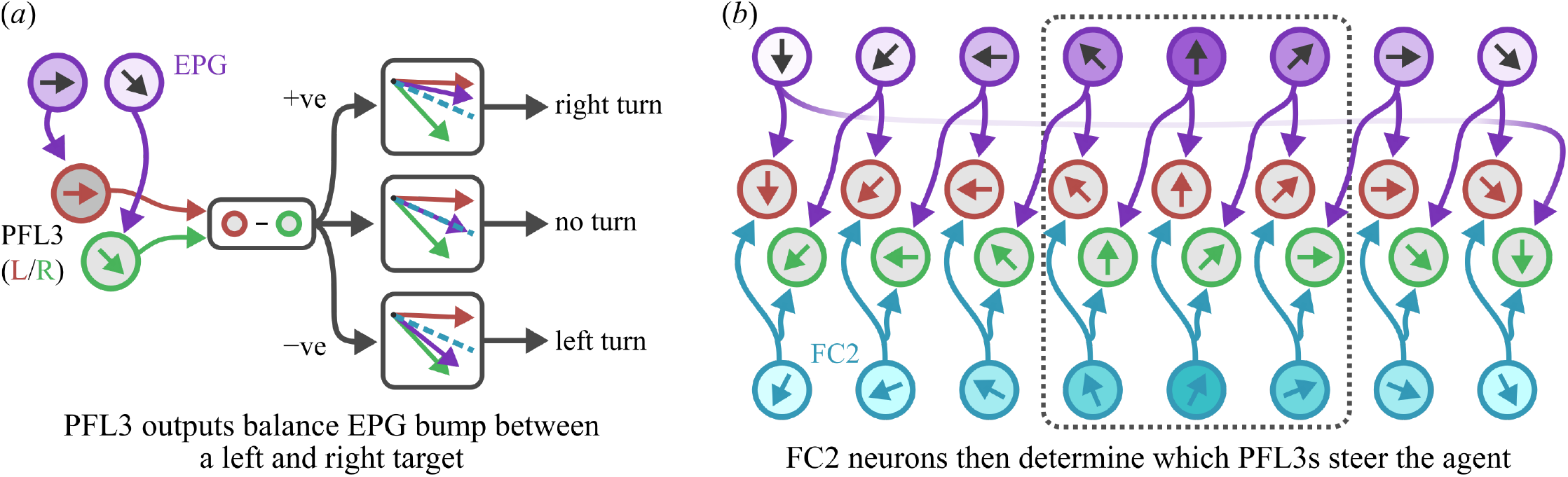
(*a*) PFL3 neuron steering mechanism. PFL3 neurons sample from neighbouring EPG neurons which creates an activity imbalance when the peak of the EPG bump does not lie between the preferred directions of those EPGs. This imbalance can be used as a steering signal, but an additional component is required to select the specific PFL3s used for steering. (*b*) In order to steer along a specific bearing, FC2 neurons scale PFL3 activity so that only specific PFL3s contribute to steering output. Note that the goal direction represented by a given FC2 neuron is defined by the PFL3s that neuron innervates (which in turn is inherited from the EPG sampling). The circuit drawn here is the one used for our experiments.

The input to PFL3 neurons is given by:

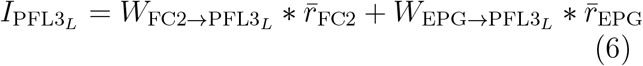

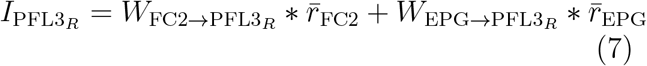

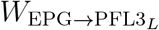 and 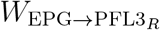 are offset from each other by a single column. By scaling the weight matrices, we can set the relative inputs from the EPGs and FC2s.

A final steering signal is generated by summing the total activity on the left and right and taking the difference [36] (also see Stone et al. [45]). The activity imbalance generates a left or a right turn:

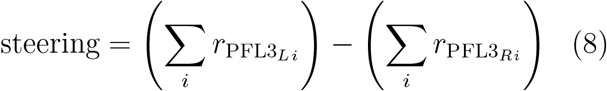

Left/right PFL3 activity will be balanced when the robot is pointing towards its goal direction, and maximally unbalanced when the robot is pointing *±*90^*°*^ from its goal direction. Note the PFL3 activity will also be balanced when the robot is pointing 180^*°*^ from its goal direction. This problem has been described as ‘false nulling’ and it has been suggested that PFL2 neurons correct for this [50]. However we did not find it necessary to include PFL2 neurons in the current model, as we observed that the ‘false null’ state is inherently unstable as any deviation will unbalance the PFL3 neurons and steer the robot back to the goal direction.

### 2.3 Robot experiments

All robot experiments were conducted at the Department of Biology, Lund University, Sweden during September 2023.

#### 2.3.1 Platform

The robot was constructed using a TurtleBot3 (Burger) kit from 2019, based on the Raspberry Pi 3B+. The platform comes with built-in motor encoders and an inertial measurement unit (IMU). The robot was additionally fitted with a camera (B0103, Arducam) with wide-angle lens, which was positioned facing upwards to detect sky cues, as well as a custom designed wind sensor.

The wind sensor was of a typical vane and anemometer design. Each component was based on the AS5600 magnetic encoder (AMS-Osram, packaged by Seeed studio) which detects the relative angle of a nearby magnet. These were used either to detect the angle of the vane (relative angle to the wind) or the speed at which the anemometer was spinning (wind speed). The wind sensor is shown in figure 3a. An I2C multiplexer (TCA9548A; Texas Instruments, packaged by Adafruit) with a custom breakout board was also added to allow the Raspberry Pi to communicate with the magnetic encoders. The angles detected by both sensor systems (angle to the bright light and angle to the wind) were used as input to the neural model (see §2.3.6).

**Figure 3.**
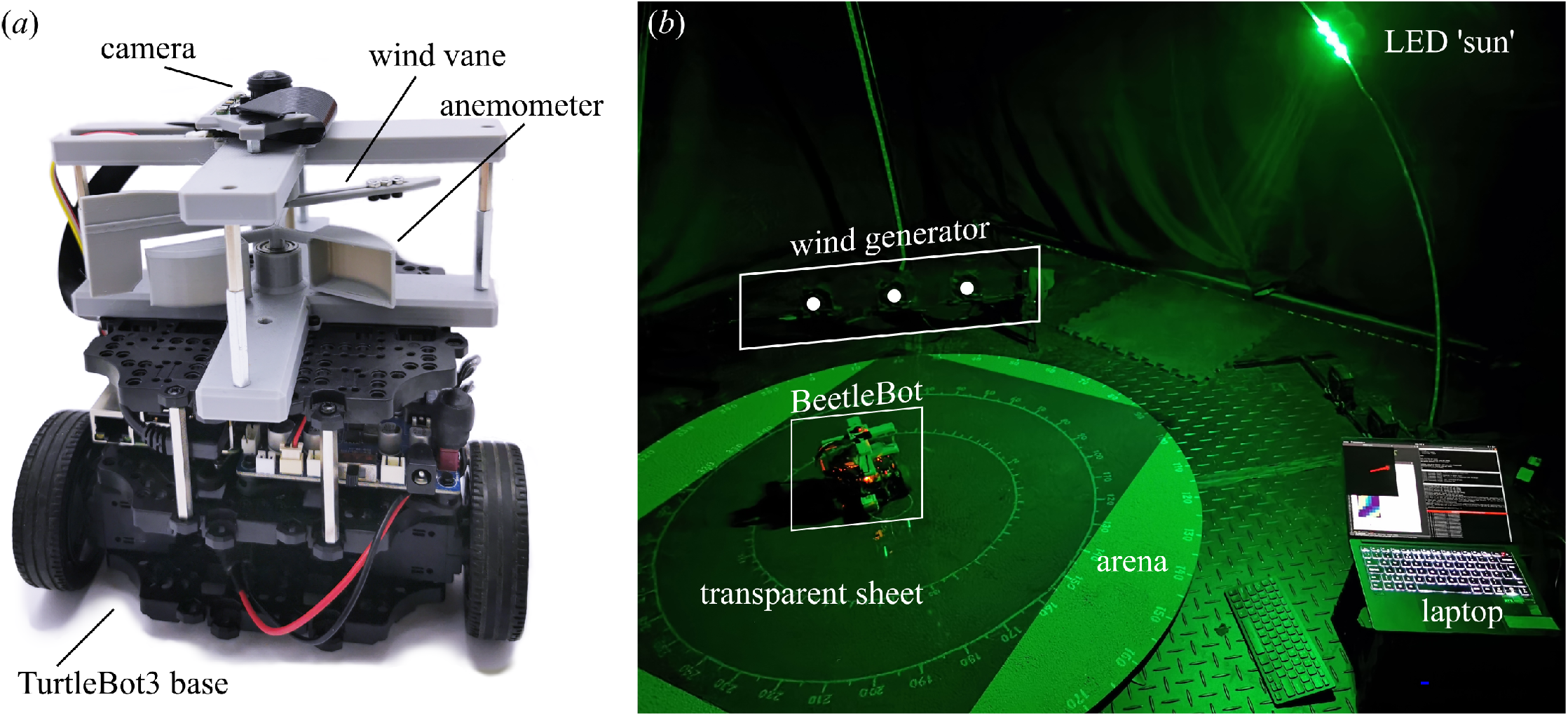
(*a*) The robot. The TurtleBot base has been consolidated to reduce the height of the robot and make space for the wind sensor. Wind direction is measured using the vane which hangs from above and speed is measured using the anemometer mounted below. (*b*) The robot in the experimental arena.

#### 2.3.2 Experimental Arena

The experimental setup (figure 3b) is similar to that used in recent dung beetle experiments [44] with some minor adjustments made for the robot. The setup is comparable to that used by Dacke et al. [10].

A circular arena with a textured surface (*r* = 0.5*m*) and edges marked in 5^*°*^ increments was placed beneath two crossed metal arches (*r* = 1.5*m*). These arches were lined with LED strips (Dotstar; Adafruit industries, New York USA). Clusters of three LEDs were used to generate an ersatz sun cue. The stimulus pattern and intensity were chosen so that the robot could reliably detect the light cue if it was in view. A transparent plastic sheet was placed over the textured arena surface so that the robot could move smoothly, while still allowing the angular increments to be read. Reflections from this sheet were not visible to the robot. The setup was configured to provide an ersatz sun cue at approximately 45^*°*^ elevation in two possible locations (180^*°*^ apart in azimuth). The robot was operated from a laptop within the tent environment. Light from the laptop was not visible to the robot.

A wind cue was provided by a custom-built wind generator placed 0.9*m* from the arena centre. The generator consisted of three fans (PFR0912XHEE, 4.50 A; Delta Electronics Inc., Taipei City, Taiwan) distributed evenly over 1*m*. Wind speed and (radial) generator position were chosen so that the robot could detect the air current when on the opposite side of the arena. The wind generator was placed such that it could be *±*90^*°*^ azimuthally offset from a given sun cue.

#### 2.3.3 Robot handling and behaviour

The robot was placed in the centre of the arena and the neural model was initialised. On command, the robot would rotate through 360^*°*^ on the spot (mimicking dung beetle ‘dance’ behaviour [11]) controlled by the on-board IMU. It would then select a random goal direction, which was encoded in the modelled FC2 neurons. The robot would then attempt to move in this goal direction until it reached the edge of the arena. The robot stopped when it reached a displacement of 0.5*m* from its start point (dead reckoning with inputs from the IMU and motor encoders). The exit angle was noted and the robot was placed back in the centre of the arena.

#### 2.3.4 Experimental procedure

Each robot trial corresponded to an individual beetle trial and involved exiting the arena four times with the neural model (and any adaptations that accrued) persisting over all four exits. On the first exit, the robot had a single initial cue available (wind or sun). On the subsequent two exits, the robot had two cues available (wind and sun). On the final exit, the initial cue was removed. For each exit, the robot’s initial orientation was randomly chosen to ensure that the model was actually trying to steer along a set direction. After the fourth exit, the neural model on the robot was re-initialised and this was treated as a new individual.

The choice of initial cue was varied and the azimuthal offset between the cues was varied such that it was either 90^*°*^ or *™* 90^*°*^. This was done to ensure that the choice of initial cue or offset direction did not bias our results. Wind-only conditions also included an ersatz sun (cluster of three LEDs) present in the zenith, to mimic previous beetle experiments [10, 44, 35].

#### 2.3.5 Robot control

During menotaxis, steering was controlled by a proportional derivative (PD) controller which operated on differential steering neuron output (equation 8). Large differences in steering neuron output were interpreted as large orientation errors. This is sufficiently true to be useful, but is susceptible to ‘false nulling’ [50] (though in practice this was not a problem, see above).

Where orientation errors were very large the robot would halt and correct on the spot. Forward movement would resume once the orientation error was reduced below a threshold. This behaviour was chosen to reduce the effect of the control scheme on the results.

#### 2.3.6 Cue detection

Cues were simply treated as either present or absent; wind speed or perceived light elevation did not alter the weight given to a cue (though see [44] and [35]). For all data in figure 4, cue weight was normalised such that single cues had a weight of 1; if both cues were present, each had a weight of 0.5 (cue ‘weight’ refers to the amplitude of the sinusoid encoding a given cue in the ER neurons, see [35]). For figure 5b and c, this normalisation was disabled to see if it had any effect on behaviour.

**Figure 4.**
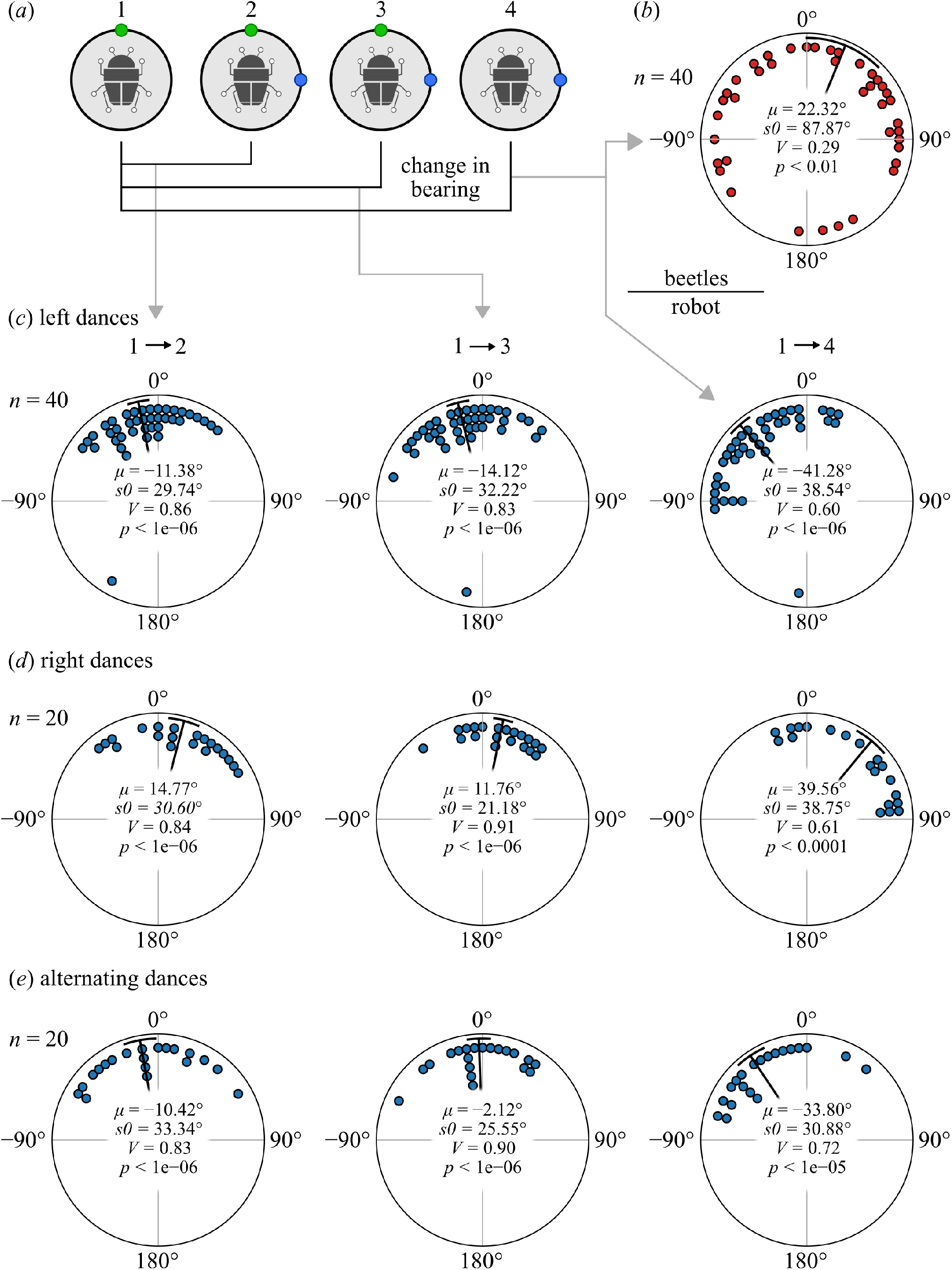
(*a*) The assay. Each individual performs one exit with an initial cue, then two with both cues, and a final exit with the initial cue removed. Cue separation was either *±*90^*°*^ and the choice of initial cue was alternated (all data are combined for analysis). (*b*) The dung beetle data from Dacke et al. [10] (reproduced with permission). (*c*) The robot data where all individuals danced with a left rotation. (*d* ) Rightward dances. (*e*) Alternating dances (R, L, R, L). For the beetles, only the first and fourth exits were compared. For the robot, comparisons are included for the first and all subsequent exits.

**Figure 5.**
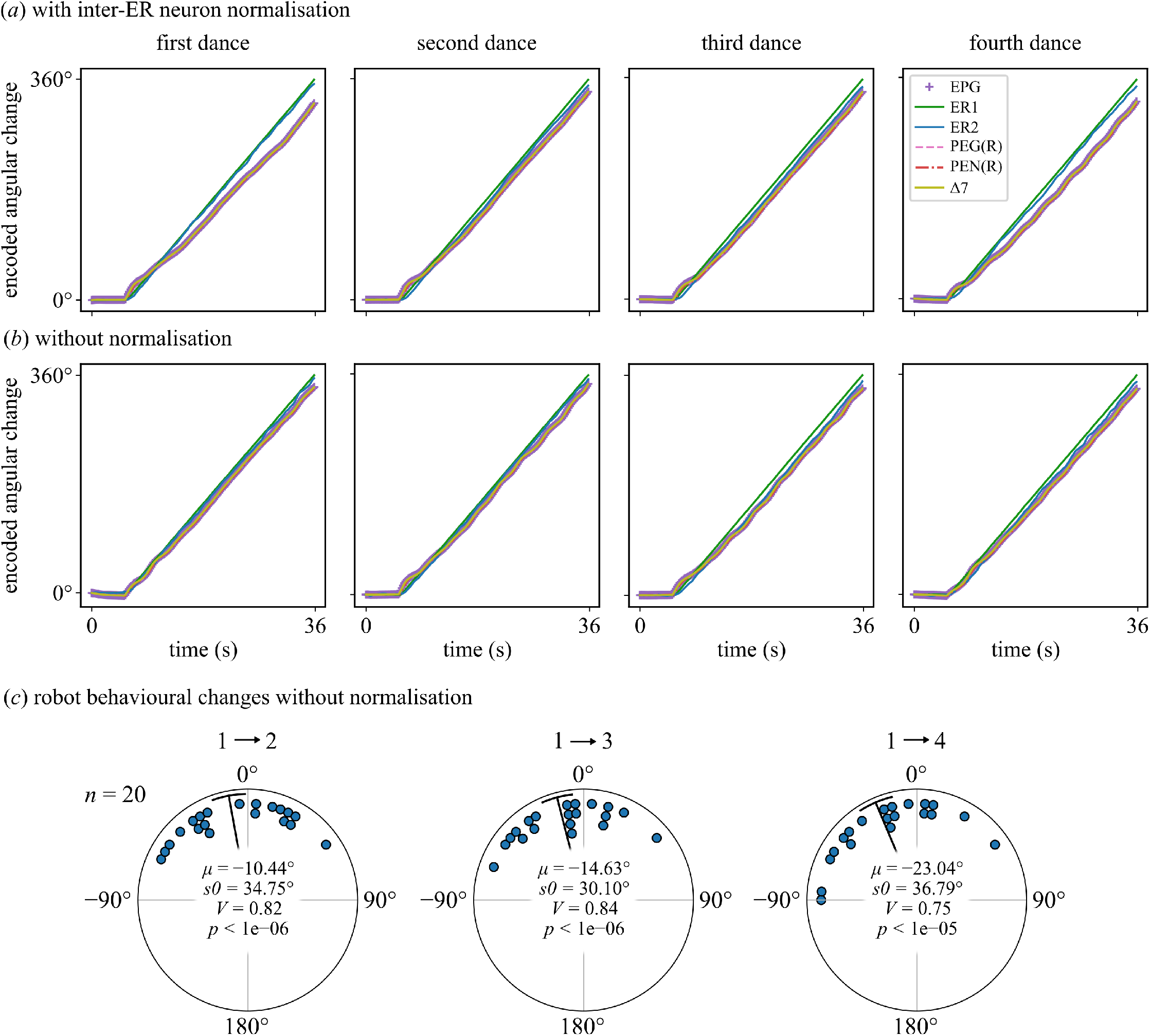
Angle encoded by each neural population relative to starting position over the course of each dance (averaged across the population). (*a*) Left-dancing individuals with inter-ER neuron normalisation enabled. In all dances there is a gradually increasing delay in neural populations downstream of the ER neurons, and this is larger in the first and fourth dances, which occurred with single cues. (*b*) Left-dancing individuals without inter-ER neuron normalisation. The increase in the delay is similar for all dances. (*c*) The robot behavioural changes (as in figure 4) where normalisation is removed, showing a decrease in the bias on the final exit.

Any rotation of the anemometer was taken to indicate wind presence and wind direction was then determined from the vane. The ersatz sun was detected by identifying the brightest spot in the image and drawing a vector from the image centre to this spot (the ‘brightest vector’). The length of this vector indicates perceived elevation and was thresholded so that only lights below a perceived elevation of approximately 70^*°*^ were detected [44, 35].

This meant that, for wind-only stimuli, the light in the zenith had no effect (though occasionally it was detected when the robot was at the arena’s edge). Practically, presence of a dominant light source ensured that no visual noise was detected as an erroneous brightest point. The light in zenith is required for motivation in dung beetles and is assumed not to affect their behaviour. Given that dung beetle orientation precision drops sharply at high light elevations [44], it is likely that they cannot use the zenith light for orientation (though this geometrically depends on the radius of the light arch and radius of the arena).

#### 2.3.7 Analysis

Circular statistical analysis was conducted using custom-made python software [35]. Each individual’s change in bearing between the first and each subsequent exit was collected, and the set of all changes was compared to a predicted mean of 0^*°*^ using the V test [10, 3]. The comparison between the first and fourth exits specifically replicates the data reported by Dacke et al. [10]. The population mean vector is computed then projected onto a unit vector pointing in a pre-chosen predicted direction. The test statistic *V* ranges from *™* 1 to 1, where values approaching 1 indicate that the population aligned with the prediction and approaching *™* 1 indicate the population was orientated in the opposite direction.

Model neuron activity was recorded throughout the experiment and analysed offline. We used this data to analyse the angle encoded by each neural population over the course of each dance period (figure 5a and b). The angular trace for each individual was shifted so that it began at zero, then we took the circular mean [3] over all individuals.

## 3 Results

### 3.1 The model replicates beetle behaviour

The robot was tested in the modality transfer experiment paradigm shown in figure 4a. It first rotated and picked a random direction to exit the arena with only one cue present (wind or light were alternately used as the first cue). It then twice experienced both cues, offset by either *±* 90^*°*^, rotating and attempting to exit the arena in the same direction as before. Finally it experienced only the second cue, again rotating and exiting the arena. Throughout the entire trial (all rotations and exit paths) the ER to EPG connections are plastic as described in the methods; with both learning rate and regulation of ER influence dependent on angular velocity.

Our results are shown in figure 4c, with the equivalent dung beetle experiment in figure 4b (data from Dacke et al. [10]). The beetle data shows the difference in direction between the first and fourth exit, and though noisy, shows statistically significant correspondence to the predicted mean difference of 0^*°*^. We plot the robot data for the difference between 1st and 2nd, 1st and 3rd, and 1st and 4th exits, and find in each case the data meet the same statistical criterion. Specifically, the results for the 1st to 4th comparison show the robot can use the second directional cue on its own to maintain the heading originally chosen relative to the first directional cue, demonstrating cue calibration (or modality transfer).

### 3.2 Dance direction biases behaviour

While the mean change in bearing is statistically near the predicted 0^*°*^, the relatively tight clustering of the robot data suggests the observed deviation from that prediction is systematic. Note this does not relate to the left or right offset of the second cue from the first, which was varied across the trials. Rather, it appears to depend on the direction in which the robot rotated before making the exit. In our first set of trials, which uniformly used left (anticlockwise) rotation, the final direction taken was systematically to the left of the expected direction (figure 4c). Repeating the experiment using only right (clockwise) rotations resulted in a clear rightward bias (figure 4d). We repeated the experiment again with alternating dances (right, left, right, left) to see if this cancelled out the effect (figure 4e) but still observe substantial leftward bias in the final exit, following a leftward rotation. We note that for the beetles, the direction of rotation of their dances was not recorded, so it is unclear whether this effect is present in the beetle population tested in Dacke et al. [10].

### 3.3 Bias results from neural lags, compounded by learning

The observed behavioural bias is indicative of a potential mismatch between the actual heading of the robot and the direction indicated by the EPG bump. One inherent source for this mismatch is the temporal difference between EPG inputs: the EPG bump is influenced by immediate sensory input (external cues and self rotation) but also by recurrent feedback from PEGs and PENs (past network state). Effectively, one set of inputs is trying to update the bump and the other is trying to keep it where it is. As the robot rotates the bump will tend to lag slightly behind the true orientation, creating an offset.

During random or corrective (oscillatory) movements, the direction of this offset will change (which may effectively form a low-pass filter on rotational inputs). But during sustained rotation in a single direction, this offset will stay consistent. This creates the potential for the adaptation process to start mapping the external cue(s) to the compass with this offset. The remapping then creates a positive feedback loop: the mapped offset will compound the inherent delay, this increased offset will trigger further remapping, and so on, continuously increasing the offset for as long as the sustained rotation continues.

This process can be seen by plotting the angles encoded by each layer of the head direction model over the course of the dance (averaged over the population). For the left-dancing individuals in figure 4c, these are plotted in figure 5a. It can be seen that the EPGs (and all ‘downstream’ populations) drift on every dance, that is, the direction indicated by the bump gradually falls further behind the direction of the external encoded cues (ER1 and ER2). The effect is more pronounced on the first and fourth dances where only a single cue was present. We hypothesised that this was a consequence of the normalisation of ER neuron population activities which, as implemented here, results in higher ER neuron activity in the presence of a single cue. This results both in increased input to the EPGs and in increased plasticity, accelerating the positive feedback effect. We tested this hypothesis by removing inter-ER group normalisation and re-running the modality transfer experiment. The angles encoded over the course of each dance are shown in figure 5b. There is a noticeable reduction in the rate of drift between ER inputs and EPG encoding during the first and fourth dances (with single cues). The reduction in drift rate is reflected in the behavioural result where we observe a reduced behavioural deviation between the first and fourth exits (figure 5c).

Given this normalisation seems to cause instability in the circuit, it is worth commenting on why it was included in the model. The head direction model in use was originally proposed as a model for cue integration [35] and in this model, the amplitude of the ER signal was used to represent the abstract concept of cue ‘weight’ (see [35] for further discussion). In the context of cue integration, weights generally sum to 1, so we assumed that these populations should be normalised (and there are some neural connections which could partially implement this [35, 31]). The model should also account for cue calibration to avoid conflicting inputs but assumptions made in service of cue integration clearly impacted the model’s ability to perform accurate calibration. This is discussed further below (§4.2).

## 4 Discussion

Previous work has demonstrated that dung beetles can calibrate the various inputs to their compass, in order to navigate consistently if some cues become unavailable. This behavioural result fits neatly into the popular conceptual model for insect compass cue calibration in the central complex, where different cue modalities arrive via ER neurons and calibration is facilitated by plastic connections between ER and EPG (“compass”) neurons [38, 21, 51]. A recent study in fruit flies appears to support the model [2]. In this work, we have demonstrated that our implementation of this multimodal head direction circuit [35] can account for the observed dung beetle behaviour in the real world, by running our model on a robot which was tested using the same behavioural setup and assay as the beetles.

Here we have replicated one of the cue calibration experiments from Dacke et al. [10], specifically, using an ersatz sun and wind as the two cues. The authors performed the same assay for green and UV point sources, positioned 180^*°*^ apart (an identical experiment is also given by el Jundi et al. [17]). el Jundi et al. [20] similarly demonstrate that dung beetles are able to learn their bearing with respect to all cues present (the authors used green and UV point sources, and UV polarised light). Even if the beetles appeared to depend most on a ‘primary cue’, they were able to keep their bearing when this cue was removed. Each of these cases can be explained by our model, assuming there exists an ER population with plastic synapses for each cue modality.

el Jundi et al. [20] suggest that beetles ‘may use a memory-dependent strategy in which they learn the spatial distribution of available celestial cues’. They call this memory a ‘snapshot’ in keeping with earlier work [1]. The snapshot concept suggests that the beetles are doing a form of visual matching [6] to perform straight-line orientation, in which the animal records the spatial distribution of available cues with respect to the goal direction when it is selected, implicitly calibrating them with each other. The alternative presented here, that the spatial distribution [20] is learnt in the ER *→* EPG neuron substrate during rotational movements, aligns with the observation that dung beetles appear to learn the cue distribution while dancing (i.e. while rotating) [20]. It also suggests the calibration precedes and is independent of selecting a goal direction. Hence, the goal direction may be stored in a different neural substrate; here we use FC2 neurons, but PFN/CPU4 neurons have also been suggested [19].

### 4.1 Cue calibration in other insects

The central complex circuitry is highly conserved across insects and it might be expected that many other insect species face behavioural situations in which calibration of different compass cues would enhance behaviour. Here we discuss several examples, but note that few provide unambiguous evidence that the animal can learn arbitrary relations between directional cues that allows one to be substituted for one another in guiding directed movements.

Bushcrickets have been shown to associate (the direction indicated by) a mating call with a visual scene, allowing a male cricket to maintain its direction toward a hidden female, even if she stops singing [48]. This could be explained by our circuit if we assume auditory information follows a pathway into ER neurons, but no such pathway has been reported to date. Our model would also require significant rotational movements in the presence of the auditory cue to establish the relationship (not reported by Von Helversen and Wendler [48]). In this case, the behaviour might be better explained by the courtship song setting a goal direction that the animal can then maintain with respect to a visual compass.

Desert ants appear to perform compass cue calibration when they first leave the nest [28]. During their transition from underground worker to overground forager, the ants will perform well-characterised learning walks outside the nest [23]. These walks are guided by geomagnetic cues initially, but over time, celestial cues will come to dominate during foraging behaviour [27, 24, 28]. This could be explained by the ant learning an ER *→* EPG mapping for skylight information, with respect to an existing map for magnetic information, consistent with the model presented here. Magnetic information is thought to be detected via the Johnston’s organ in ants so it is reasonable to suppose that magnetic information reaches the ring neurons via a similar pathway to wind [38].

Dependence on a celestial compass for navigation introduces a problem for a foraging ant: the movement of the sun over the course of the day. Potentially ER*→* EPG remappings could solve this, without requiring an internal clock for time compensation, if each time the ant leaves the nest she remaps celestial information according to a (relatively) stable magnetic reference. However, this would require modification of our current model, in which there is no such thing as a primary or reference cue, only stronger or weaker connections formed by experience. In this case, experience would lead the sun cue to dominate, causing other cues to be remapped according to it, rather than vice-versa. Alternatively, if celestial inputs were time compensated before reaching the ER neurons (as in a recently proposed circuit model [25]), then from the perspective of the circuit, the relationship between solar and magnetic information would appear to stay constant, and any remapping would simply reinforce the relationship. Our model can explain an initial calibration, but time compensation seems better suited to maintenance.

Additional work in desert ants has reported a transfer of information between different compass cue modalities [33]. In these experiments, the ants were trained using either the sun or polarised skylight in isolation, and observed to orientate consistently if presented with only the untrained cue. It is unclear from these experiments if the ants learned the relationship between polarised light and solar position prior to experimental training (consistent with our model) or if they innately expect these cues to exhibit their natural spatial relationship. Unlike some dung beetle experiments [20], there was no attempt to train them with cues that conflicted with the natural relation of sun position and polarisation pattern. In similar experiments on honeybees, it has generally been assumed that the relationship of cues is innately encoded. Rossel and Wehner [41] demonstrate that honeybees appear to expect a specific relationship between solar position and the atmospheric polarisation pattern. This expectation also applies to the spectral content of the sky. Honeybees have been described as interpreting (unpolarised) UV light as if it were an ‘anti-sun’ cue [15, 4, 41]. It has been suggested that the bees use a filter matched to the spectral content of the sky (high amounts of green in the solar hemisphere, high amounts of UV in the anti-solar hemisphere) [20]. Rossel and Wehner [41] describe the bees as having an ‘internal map of the sky’, and Brines and Gould [4] describe them as having a ‘sun-sky rule’. However, as in the ant case, the data do not allow us to distinguish whether this rule is learned or innate. These experiments were conducted by training the bees to a feeder, then studying their dances under artificial stimuli to determine how they interpreted the different wavelengths (or polarisation patterns [41]). Therefore, it is possible that the bees learned their ‘map of the sky’ during foraging, in much the same way that the dung beetles learn the spatial relationship of cues.

### 4.2 Additional insights into CX plasticity

Providing an implementation of the conceptual model given in the literature [38, 21, 51], allows for a critical examination of the proposed system. While our implementation successfully captures the target behaviour, several issues arose which are relevant to future exploration of this circuit.

The model experiences some systematic error in the form of a compass lag induced by the time difference between instantaneous and recurrent inputs. Similar compass lag could exist in dung beetles, but be hidden in population data if dance directions were evenly split. Dirlik [13, Paper I] provides a systematic investigation of dung beetle dance directions, influencing factors, and resultant influences on behaviour. The authors note that *Kheper lamarcki* individuals experience a small bias in their travel direction which is influenced by dance direction. Dancing counter clockwise resulted in a slight leftward shift and vice versa, matching the effect we observed with the robot (though with smaller magnitude). This suggests that the real circuit may have a mechanism to compensate for the compass delay and remapping cycle. Such a mechanism could be behavioural (compensatory movement patterns) or neurophysiological.

In fruit flies, Seelig and Jayaraman [43] note that the EPG bump did not always respond instantly to cue movement, which could be symptomatic of the effect we observed in the model. A more recent study corroborates this delay in the EPGs, and explores the neurophysiology underlying the associative plasticity mechanism in the circuit [40]. The study suggests that associative learning is mediated by EL neurons which receive input from EPGs and connect back to ER neuron axons across the Ellipsoid Body. EL outputs are octopaminergic, and it is proposed that ER to EPG synaptic strength (inhibitory) is depressed where there is coincident activation of ER and EL neurons. This mechanism could be represented by the abstracted learning rule we have used in the current study (equation 2): if EL activity is assumed to be driven directly by EPGs, then substituting EL for EPG activity in this equation would effectively be an identical model. However, although the EL neurons, like the EPGs, exhibit an activity bump which follows the direction of the animal, the EL bump appears to precede the EPG bump during rotations [40], possibly due to the additional input they receive from PENs. This means the learning signal from the EL does not suffer from the problematic delay in EPG updating that produced a behaviour-dependent drift in the association between cues in our study. The functional characteristics of EL neurons and their role in learning was not known when our model was constructed, but could be integrated into future work to see if this drift can indeed be removed.

We also found the circuit very sensitive to changes in the relative strength of different inputs (e.g. sensitivity to normalisation, figure 5). Tuning the recurrent circuitry to operate as a ring attractor is a delicate balancing act in itself [37]. Introducing plastic connections for calibration adds the requirement that the recurrent circuitry remain stable regardless of the status of those connections (and the amount of input they provide to the circuit). This creates a problem when trying to learn the spatial relationships of different cues. For the learning mechanism to work, PENs must accurately communicate angular velocity to the EPGs. But the relative influence of PENs over EPGs changes as ER *→* EPG connections are updated. In the literature, the interaction between these neural populations seems to be treated as a solved problem. However, other computational models have either omitted plastic inputs [32, 39, 26, 46], or omitted ER neuron input during learning [8, 12]. Understanding how relative inputs from different neuron classes are regulated as the connection strengths change constitutes an interesting line for future investigation.

This issue arose because we required the robot model to operate consistently during learning and navigating, rather than isolating these functions for modelling. Similarly, we found that previous assumptions made when focussing only on how the circuit could perform cue integration (e.g. inter-ER neuron normalisation) were detrimental to its ability to perform accurate cue calibration (figure 5). This highlights a larger issue in circuit modelling, where too much focus can be given to solving individual tasks and insufficient consideration given to other roles the same circuit needs to perform. While functional compartmentalisation is required to some degree for pragmatic model construction and validation, we believe there is scope for more model integration than there has been in the past. Even if unsuccessful, attempted integration would still be informative as it can point out the shortcomings in our models. Similar utility can be gained by taking models out of simulation and providing robotic implementations.

### 4.3 The benefits of robot testing

The experiment presented here highlights why robot testing is useful in a literature dominated by computer simulation. Implementing this work on a physical robot led to the discovery of a few shortcomings of the simulation used by Mitchell et al. [35].

As an example, wind was a far more reliable cue than was assumed in that work, and moreover, it is clear that the error in perceived wind position was not evenly distributed about its true position (i.e. von Mises with zero mean as assumed in [35]). If anything, the wind input lags ever so slightly compared to the intensity cue (figures 5a and b). This implies that the ER*→* EPG mappings formed by wind would perhaps be slightly skewed, but they would not experience significant disruption. In turn, this makes it extremely unlikely that disruption to the ER *→* EPG mapping would actually affect cue influence (i.e. that the connections would encode cue reliability in a way which significantly impacts behaviour [51], also see [35]). This conclusion is obviously influenced by sensor design. But this exposes the fact that simulating noisy wind detection using a von Mises process neglected any consideration of its biological realism compared to a bilateral antennal array [47]. The robot makes such assumptions clear by having a real sensor instead of an abstract representation.

The simulation also included a simplistic notion of time as it was assumed that simulation time could be scaled in order to operate on an arbitrary scale, which was not true. For example, a small amount of PEN activation over a long time frame is not the same as a large amount of PEN activation over a short time frame. Low PEN activation can fail to overcome recurrent input from the PEGs and fail to update the compass at all. This only became apparent when the model was operating on the robot in real time.

In simulation, even a stochastic sequence of events is quite tightly controlled. As soon as the model was running on a robot which was being handled in an arena, it became clear just how unintentionally sterile the simulation was. With the benefit of hindsight it is easy to argue that sensing and time could have been better represented, and some kind of handling model could have been included. The broader take-home is that there are many things we *could* account for in simulation which a robot gives us for free [49]. We are not arguing that simulations are useless; on the contrary, they serve as a necessary starting point. Robotic implementations undoubtedly have their problems, but these are usually obvious from the design of the system. The robot is forced to operate in the real world, exposing gaps in our understanding of the computational sequence from perception to behaviour.

## 5 Acknowledgements

From the Lund Vision Group, we would like to thank Shahrzad Shaverdian, Yakir Gagnon, and Elin Dirlik for their help with the behavioural setup. Additional thanks go to Elin Dirlik for assisting with photo and video collection. We would also like to thank all those in the Dacke Lab who spent days ‘on-call’ as experimental assistants. Finally, we would like to thank the members of the InsectRobotics group for their feedback on the reporting phase (with particular thanks to Evripidis Gkanias for his input).

## 6 Code and data availability

Our model code, analysis code, and raw data are available at https://doi.org/10.5281/zenodo.15001500.

## 8 Author Contributions

RM: Conceptualisation, Investigation, Methodology, Software, Visualisation, Writing - original draft, Writing - review and editing. MD: Conceptualisation, Funding acquisition, Project administration, Resources, Supervision, Writing - review and editing. BW: Conceptualisation, Funding acquisition, Project administration, Resources, Supervision, Writing - review and editing.

