## Supplementary information for "A robot model of compass cue calibration in the insect brain"

### 1 Model specification

The majority of the neural model is carried over from Mitchell et al. [2], therefore much of this specification is identical to that work.

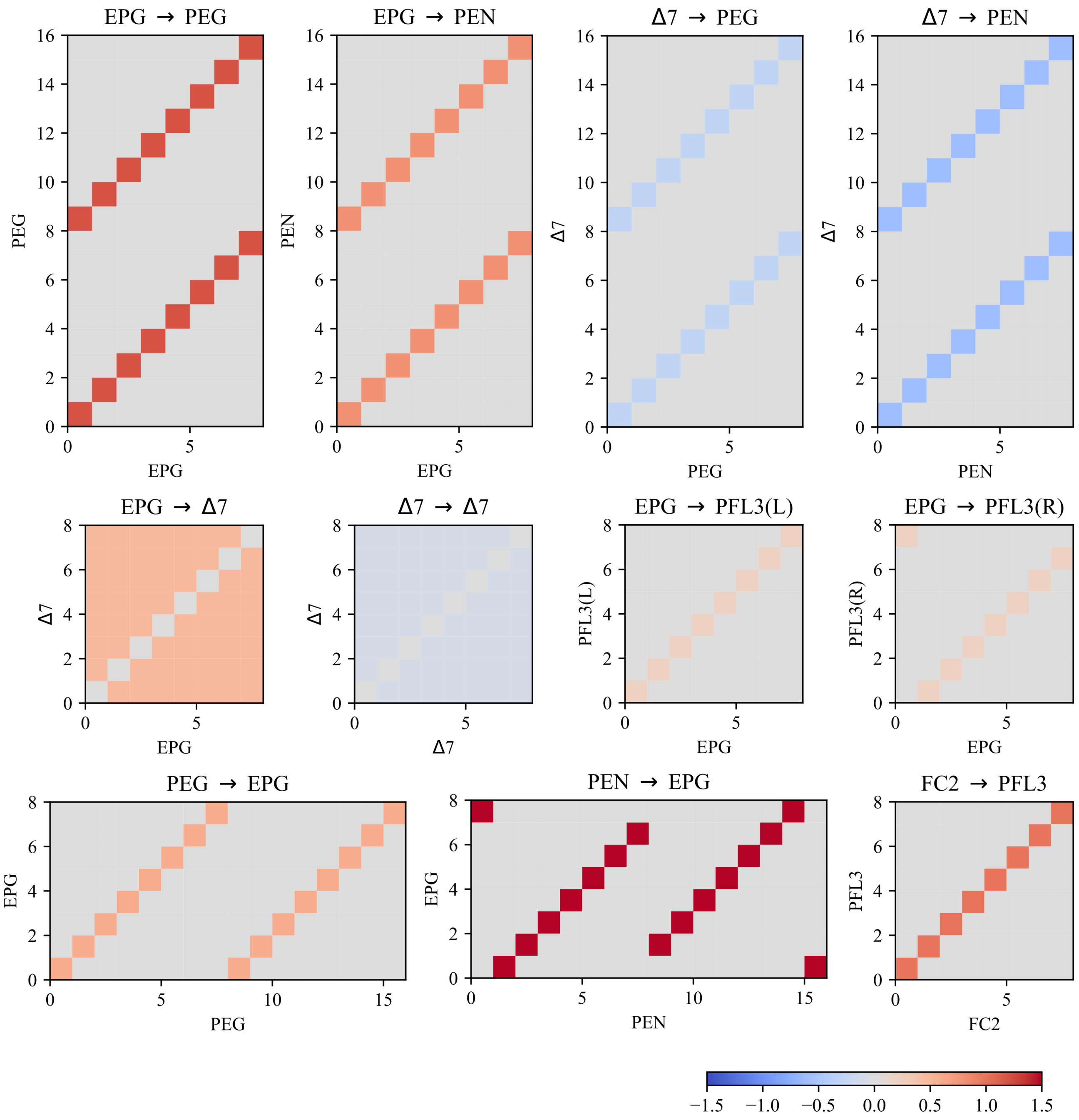


Figure 1: Adjacency matrices documenting neuron interactions in the model. Inhibitory connections are blue, excitatory are red. An absence of connection is grey. ER → E-PG connections are omitted. These represent a slightly re-tuned version of the network from [2], with added PFL3 and FC2 neurons (and relevant connections). PFL3/FC2 wiring is bio-inspired but not anatomical, for justification see [1].

#### 1.1 Noisy firing rate model

We use the same firing rate model as Mitchell et al. [2], but with added noise. Firing rate $r$ is given by

$$r = \frac{1}{1 + e^{-(aI+b)}} + \epsilon \tag{1}$$

where $I$ is the input, $a$ and $b$ give the slope and bias of the activation function, and $\epsilon \sim N(0, 0.001)$. We denote individual firing rate as $r$ and use $\bar{r}$ to denote a vector of firing rates across a population. Parameters $a$ and $b$ were tuned by hand for each neural population. Parameters for different neuron types are given in Table S1.

Table S1: The neuron count $n$ and sigmoid tuning parameters used for each neuron type, corresponding to $a$ and $b$ in Equation 1. There are 8 ER neurons in each group, and PFL3 neurons split into equally-sized L and R sub-populations.

| **Neuron class** | $n$ | $a$ | $b$ |
|---|---|---|---|
| ER | 8 | 4 | 1.8 |
| EPG | 8 | 4 | 1.8 |
| PEN | 16 | 4 | 5 |
| PEG | 16 | 4 | 3 |
| Δ7 | 8 | 3 | 3 |
| FC2 | 8 | 4 | 1 |
| PFL3 | 16 | 4 | 3 |

#### 1.2 Neuron specifications

##### 1.2.1 ER neurons

Each ER neuron has a receptive field centred on its preferred direction $\theta_{\text{ER}} \in \{0, \frac{\pi}{4}, \frac{\pi}{2}, \frac{3\pi}{4}, \pi, \frac{5\pi}{4}, \frac{3\pi}{2}, \frac{7\pi}{4}\}$. The ER neuron input is

$$I_{\text{ER}_{l,i}}(t) = \frac{\cos(\theta_{\text{ER}_{l,i}} - \theta_C)}{2} + 1 \tag{2}$$

(adapted from Stone et al. [4]), where $\text{ER}_{l,i}$ is the $i$th ER neuron of group $l$, $\theta_C$ is the azimuthal angle of the input cue, and $\theta_{\text{ER}_{l,i}}$ is the preferred direction of that ER neuron.

**ER sinusoidal amplitude and normalisation** Sinusoidal amplitudes for $\text{ER}_1$ and $\text{ER}_2$ are governed by parameters `w1` and `w2` respectively. These amplitude modifiers are applied after the ER rates are computed (i.e. we scale the ER responses):

$$\bar{r}_{\text{ER}_l} := \mathtt{wl} \cdot \bar{r}_{\text{ER}_l} \tag{3}$$

Following Mitchell et al. [2], we assumed that $\mathtt{w1}+\mathtt{w2} = 1$ so that input to the EPGs remains within an acceptable range, regardless of which amplitude is higher. Mitchell et al. [2] made this assumption with the context that the network would be performing cue integration, where cue weights are generally normalised.

**Disabling normalisation** As noted in the manuscript, we found that normalisation disrupted the cue calibration functionality of the network; we therefore investigated the effect of removing it. This means that if a cue was detected by the robot, it received a weight of 0.5 and 0 otherwise.

##### 1.2.2 PEN neurons

PEN input is given by:

$$I_{\text{PEN, Left}}(t) = (1 - h(v_\theta)) + W_{\text{EPG}\rightarrow\text{PEN}} \cdot \bar{r}_{\text{EPG}}(t-1) + W_{\Delta7\rightarrow\text{PEN}} \cdot \bar{r}_{\Delta7} \quad (4)$$

$$I_{\text{PEN, Right}}(t) = h(v_\theta) + W_{\text{EPG}\rightarrow\text{PEN}} \cdot \bar{r}_{\text{EPG}}(t-1) + W_{\Delta7\rightarrow\text{PEN}} \cdot \bar{r}_{\Delta7} \quad (5)$$

With $v_\theta$ being the signed angular velocity in $°/0.2s$. The terms $h(v_\theta)$ and $(1 - h(v_\theta))$ above generate the differential speed encoding across the P-EN population while keeping overall activity constant (if one side's average activity increases, the other's decreases).

The function $h$ is given by:

$$h(x) = [k \cdot x + 0.5]_{[0,1]} \quad (6)$$

With $k = 0.5/20.9$, and $[x]_{[y,z]}$ denotes the clipping operation $[x]_{[y,z]} = \max(y, \min(x, z))$. The value of $k$ (tuned by hand) roughly defines PEN activity for a given angular velocity, and therefore how that angular velocity translates into a change in EPG activity. This means the maximum angular velocity which can be detected by self-motion alone is $20.9°/0.2s$, or $104.5°/s$. This maximum proved sufficient for the robot.

##### 1.2.3 EPG neurons

EPG input is given by:

$$I_{\text{EPG}}(t) = S_{\text{ER}} \cdot (W_{\text{R}_0\rightarrow\text{EPG}} \cdot \bar{r}_{\text{ER}_0}(t) + W_{\text{ER}_1\rightarrow\text{EPG}} \cdot \bar{r}_{\text{ER}_1}(t)) + W_{\text{PEN}\rightarrow\text{EPG}} \cdot \bar{r}_{\text{PEN}}(t) + W_{\text{PEG}\rightarrow\text{EPG}} \cdot \bar{r}_{\text{PEG}}(t-1) \quad (7)$$

$S_{\text{ER}}$ scales ER input to EPGs depending on angular velocity:

$$S_{\text{ER}} = -1.4 \cdot (1 - [\rho \cdot |v_\theta|]_{[0,0.8]}) \quad (8)$$

where $|v_\theta|$ is the absolute angular velocity and $\rho = 0.833$ is a tuning constant which defines how sharp the inhibition will be as angular velocity increases.

##### 1.2.4 Δ7 neurons

Δ7 input is given by:

$$I_{\Delta7}(t) = W_{\text{EPG}\rightarrow\Delta7} \cdot \bar{r}_{\text{EPG}}(t) + W_{\Delta7\rightarrow\Delta7} \cdot \bar{r}_{\Delta7}(t-1) \quad (9)$$

##### 1.2.5 PEG neurons

PEG input is given by:

$$I_{\text{PEG}}(t) = W_{\text{EPG}\rightarrow\text{PEG}} \cdot \bar{r}_{\text{EPG}}(t) + W_{\Delta7\rightarrow\text{PEG}} \cdot \bar{r}_{\Delta7}(t) \quad (10)$$

##### 1.2.6 FC2 neurons

FC2 neurons have preferred directions $\theta_{\mathrm{FC2}} \in \{0, \frac{\pi}{4}, \frac{\pi}{2}, \frac{3\pi}{4}, \pi, \frac{5\pi}{4}, \frac{3\pi}{2}, \frac{7\pi}{4}\}$. Note that these preferred directions are used to generate an activity pattern but do not correspond to the actual goals represented by each neuron (see [1]).

The input for the $i$th FC2 neuron is given by:

$$I_{\mathrm{FC2}_i} = \cos(\theta_g - \theta_i) \tag{11}$$

where $\theta_g$ is the goal direction and $\theta_i$ is the preferred angle of the $i$th FC2 neuron.

##### 1.2.7 PFL3 neurons

PFL3 input is given by:

$$I_{\mathrm{PFL3}_L} = W_{\mathrm{FC2}\to\mathrm{PFL3}_L} * \bar{r}_{\mathrm{FC2}} + W_{\mathrm{EPG}\to\mathrm{PFL3}_L} * \bar{r}_{\mathrm{EPG}} \tag{12}$$

$$I_{\mathrm{PFL3}_R} = W_{\mathrm{FC2}\to\mathrm{PFL3}_R} * \bar{r}_{\mathrm{FC2}} + W_{\mathrm{EPG}\to\mathrm{PFL3}_R} * \bar{r}_{\mathrm{EPG}} \tag{13}$$

$W_{\mathrm{EPG}\to\mathrm{PFL3}_L}$ and $W_{\mathrm{EPG}\to\mathrm{PFL3}_R}$ are offset from each other by a single column (see figure 1). A final steering signal is generated by summing the total activity on the left and right and taking the difference [3] (also see Stone et al. [4]).

The activity imbalance generates a left or a right turn:

$$\text{steering} = \left(\sum_i r_{\mathrm{PFL3}_{Li}}\right) - \left(\sum_i r_{\mathrm{PFL3}_{Ri}}\right) \tag{14}$$

Left/right PFL3 activity will be balanced when the robot is pointing towards its goal direction, and maximally unbalanced when the robot is pointing $\pm 90°$ from its goal direction.

#### 1.3 Learning

The model uses the same anti-hebbian learning rule presented in [2].

$$\Delta w_{i,j} = -\eta \cdot (r_{ER_{l,j}} - \theta_{ER_l}) \cdot (r_{\mathrm{EPG}_i} - \theta_{\mathrm{EPG}}) \tag{15}$$

where $\Delta w_{i,j}$ is the change in synaptic strength from $\mathrm{ER}_{l,j}$ onto $\mathrm{EPG}_i$, and the $\theta$ terms are activity thresholds. The threshold terms will lead to a decrease in strength where one neuron is above its threshold and the other below.

The learning rate $\eta$ is proportional to angular velocity

$$\eta = \frac{c}{v_{\max}} \cdot [|v_\theta|]_{[0,v_{\max}]} \tag{16}$$

where $|v_\theta|$ is the absolute angular velocity, $c = 0.1$ is a tuning constant, and $v_{\max}$ is the maximum angular velocity which can be perceived by the network ($20.9°/0.2s$, see section on PENs).

In the previous work, learning was contextually enabled or disabled [2]. In our experiments, learning is always enabled and simply scaled depending on $v_\theta$.

**ER $\to$ EPG connection normalisation** ER $\to$ EPG connections are always normalised such that the ER *input to* a single EPG always sums to 1. This is true whenever connections are modified or in their initial randomised state.

#### 1.4 Model initialisation

In this iteration of the model, we start with a flat, noisy activity pattern in the EPGs. The noise results in slight imbalances in activity between different EPGs, and over time, the recurrent circuitry causes these imbalances to stabilise into a single bump.

No ER input is provided to the EPGs during this initialisation. We therefore require no assumptions on the state of ER $\rightarrow$ EPG mappings and start with randomised connections. This is a significant change from Mitchell et al. [2], where we required specific default connection patterns to initialise the network.
